# Conserved type VI secretion regulation in diverse *Vibrio* species by the regulatory proteins TfoX and TfoY

**DOI:** 10.1101/466458

**Authors:** Lisa C. Metzger, Noémie Matthey, Candice Stoudmann, Esther J. Collas, Melanie Blokesch

## Abstract

Bacteria of the genus *Vibrio* are common members of aquatic environments where they compete with other prokaryotes and defend themselves against grazing predators. A macromolecular protein complex called the type VI secretion system (T6SS) is used for both purposes. Previous research showed that the sole T6SS of the human pathogen *V. cholerae* is induced by extracellular (chitin) or intracellular (low c-di-GMP levels) cues and that these cues lead to distinctive signalling pathways for which the proteins TfoX and TfoY serve as master regulators. In this study, we tested whether the TfoX- and TfoY-mediated regulation of T6SS was conserved in non-cholera species, and if so, how these regulators affected the production of individual T6SSs in double-armed vibrios. We show that, alongside representative competence genes, TfoX regulates at least one T6SS in all tested *Vibrio* species. TfoY, on the other hand, fostered motility in all vibrios but had a more versatile T6SS response in that it did not foster T6SS-mediated killing in *V. fischeri* while it induced both systems in *V. alginolyticus*. Collectively, our data provide evidence that the TfoX- and TfoY-mediated signalling pathways are mostly conserved in diverse *Vibrio* species and important for signal-specific T6SS induction.

**Originality-Significance Statement:** This work provides new insight into the regulatory circuits involved in type VI secretion in diverse *Vibrio* species. Specifically, it is the first study to compare the effects of the two regulatory proteins TfoX and TfoY on the primary or secondary type VI secretion systems of non-cholera vibrios. Importantly, this work also shows that decreased c-di-GMP levels in *V. parahaemolyticus* lead to TfoY production without changing *tfoY* transcript levels, thereby indirectly linking TfoY production to surface sensing.

## Introduction

Secretion systems that enable the export of macromolecules across membranes are often associated with pathogenesis in Gram-negative bacteria. However, these machineries and their substrates also play important roles in the interactions between bacteria and other microorganisms in their natural environment (Costa *et al*., 2015; Green and Mecsas, 2016). An important secretion system, the type VI secretion system (T6SS), was first described in *Vibrio cholerae* and is a syringe-like killing device involved in virulence and inter-bacterial warfare (Pukatzki *et al*., 2006; Ho *et al*., 2014; Russell *et al*., 2014). T6SSs are present in ~25% of sequenced Gram-negative bacteria, including pathogenic and non-pathogenic species, and are defined by a set of 13 core components (Das and Chaudhuri, 2003; Bingle *et al*., 2008; Boyer *et al*., 2009). These components build a macromolecular complex that shares both structural and functional homology with contractile bacteriophage tails, such as in the phage T4 (Leiman *et al*., 2009; Basler, 2015; Taylor *et al*., 2018). Briefly, a tail-like structure, which is composed of hemolysin coregulated protein (Hcp) hexamers and sharpened by spike-like tip proteins (VgrG and PAAR), is surrounded by a contractile sheath structure and polymerizes onto a membrane-spanning multi-protein complex. When the sheath contracts, the effector-decorated tip complex is propelled outwards together with the inner tube to puncture the membrane of a neighboring target cell and inject toxic effector proteins (reviewed by (Zoued *et al*., 2014; Cianfanelli *et al*., 2016). Each T6SS encodes its own sets of variable effector proteins that determine the specificity and the biological function of the system, as they can impact both eukaryotic host cells and bacterial competitors. To prevent self-intoxication and to kin-discriminate siblings, bacteria produce effector-specific neutralizing immunity proteins, which are frequently encoded adjacent to the effector gene in a bicistronic unit (Durand *et al*., 2014; Alcoforado Diniz *et al*., 2015). Studying the various T6SS systems and their regulation is therefore important, as such knowledge sheds light onto the evolution of vibrios in response to adapting niches and their interaction with environmental hosts.

A comparison of the various T6SS-associated across *Vibrio* species indicate that there must be multiple different functions and regulatory mechanisms associated with the system(s). The structural components of the T6SS are mostly encoded within larger gene clusters, which are variable with respect to genetic content and organization. The genes coding for the secreted core components (Hcp, VgrG, PAAR) are often found in smaller auxiliary clusters together with the genes that encode the effectors-immunity pairs (Boyer *et al*., 2009). Many bacterial genomes encode more than one T6SS, which likely serve different needs (Boyer *et al*., 2009). Accordingly, bacteria employ a wide variety of regulatory mechanisms, from common cues to specific regulatory cascades, to ensure appropriate expression of T6SS genes (Miyata *et al*., 2013), which likely vary within species when multiple T6SS systems are present.

*V*. cholerae O1 El Tor strains, which are responsible for the current 7^th^ pandemic of cholera (further referred to as pandemic strains), harbor a single T6SS. In contrast, many non-cholera *Vibrio* species encode several T6SSs, such as clinical isolates of *V. parahaemolyticus* and environmental isolates of *V. vulnificus*, both of which carry two T6SSs (Yu *et al*., 2012; Church *et al*., 2016). The genome of the squid symbiont *V. fischeri*, on the other hand, codes for one T6SS even though a recent study by Speare *et al*. identified a secondary T6SS in several *V. fischeri* isolates that fostered niche domination within the squid’s light organ (Speare *et al*., 2018).

The T6SS of pandemic *V. cholerae* is silent under standard laboratory conditions (Pukatzki *et al*., 2006). In contrast to pandemic strains, the O37 serogroup strains V52 and ATCC25872 are toxigenic but non-pandemic and their T6SS is constitutively activated, as is the case for most environmental isolates (Pukatzki *et al*., 2006; Unterweger *et al*., 2012; Bernardy *et al*., 2016; Van der Henst *et al*., 2018). To explain the differences in T6SS activity and to provide insight into the T6SS’s biological functions, it is therefore important to understand the underlying regulatory pathways in pandemic *V. cholerae* strains. Accordingly, several minor and major T6SS regulators have been identified in *V. cholerae* (Joshi *et al*., 2017) by changing their abundance through deletion or forced expression of the respective genes, which resulted in changes in T6SS gene expression or T6SS activity. Two of these major regulators, TfoX and TfoY (Borgeaud *et al*., 2015; Metzger *et al*., 2016), contain TfoX-like N- and C-terminal domains, and proteins containing such domains are usually annotated as regulators of natural competence for transformation due to their presence in the competence regulator Sxy of *Haemophilus influenzae* (Redfield, 1991) and TfoX in *V. cholerae* (Meibom *et al*., 2005). We recently showed that TfoX induces the T6SS in parallel with the competence machinery at a high bacterial cell density (measured by quorum sensing [QS] and signaled through its master regulator, HapR) on chitinous surfaces, leading to T6SS-mediated killing followed by the absorption of prey-released DNA (Borgeaud *et al*., 2015). Coupling kin-discriminating neighbor killing and competence fosters horizontal gene transfer and, ultimately, evolution (Veening and Blokesch, 2017).

Interestingly, *V. cholerae* and most other members of the genus *Vibrio* (vibrios) contain a TfoX homolog named TfoY (Pollack-Berti *et al*., 2010; Metzger *et al*., 2016). However, despite the homology to TfoX and its common annotation as a competence regulator, TfoY is neither essential nor sufficient for competence induction in *V. cholerae* (Metzger *et al*., 2016). TfoY also induces the T6SS of *V. cholerae*, though the TfoX and TfoY regulons are non-overlapping, with TfoX co-inducing competence and TfoY orchestrating a defensive reaction. This defensive reaction includes enhanced motility and the co-production of the T6SS with additional extracellular enzymes, some of which are known to intoxicate amoebal predators (Metzger *et al*., 2016; Van der Henst *et al*., 2018). Moreover, TfoX is naturally produced upon growth on chitinous surfaces (Meibom *et al*., 2005), while TfoY production is inhibited by c-di-GMP, an secondary messenger in bacteria, such that it is only present at low intracellular c-di-GMP levels (Inuzuka *et al*., 2016; Metzger *et al*., 2016).

Given the presence of TfoX and TfoY in many vibrios, we aimed to understand their role in non-cholera species. We show that the T6SS activation is highly conserved across vibrios, with the majority of tested species concomitantly inducing natural competence and motility when TfoX and TfoY, respectively, are produced. Interestingly, we observed different scenarios when comparing those *Vibrio* species that encode more than one T6SS, with TfoX and TfoY not fulfilling the same inducing role for all T6SSs. We therefore conclude that some vibrios specifically induce one or the other T6SS under different environmental conditions driven by either TfoX or TfoY, while other species combine forces to simultaneously induce both T6SSs.

## Results and discussion

### Conservation of T6SS production by TfoX and TfoY in pandemic isolates of V. cholerae

Since its discovery in a non-pandemic strain of *V. cholerae*, it has been reported that the T6SS of pandemic strains is silent under standard laboratory conditions (Pukatzki *et al*., 2006). We recently showed that the two regulatory proteins TfoX and TfoY foster T6SS production and, accordingly, inter-bacterial competition in pandemic *V. cholerae* (Borgeaud *et al*., 2015; Metzger *et al*., 2016). The previous study that addressed TfoY, however, was based on a single strain of *V. cholerae*, namely strain A1552. This human isolate originated from food-borne transmission on an airplane returning to the US from Peru/South America (Blokesch, 2012a) and therefore belongs to the West-African South American (WASA) lineage (precisely, the LAT-1 sublineage) of the currently ongoing seventh cholera pandemic (Mutreja *et al*., 2011; Domman *et al*., 2017). As strain-specific differences are frequently reported, this study was designed to first confirm the generality of TfoX- and TfoY-dependent T6SS activation in pandemic strains of *V. cholerae*. To do this, we tested five isolates from three different countries (Bangladesh, Peru, and Bahrain) for T6SS-dependent killing of *Escherichia coli*. Notably, we used a derivative of the first sequenced isolates of *V. cholerae*, pandemic strain N16961 (Heidelberg *et al*., 2000), in which the frameshift mutation in the quorum-sensing regulator gene *hapR* was repaired (Kühn *et al*., 2014). As demonstrated in Figure S1, all strains behaved similarly to A1552 upon TfoX or TfoY production. We therefore suggest that the dual control over T6SS by these two regulators is conserved among pandemic *V. cholerae*.

### Functionality of TfoX and TfoY homologs from other Vibrio species in V. cholerae

Pollack-Berti and colleagues showed that TfoX and TfoY existed in all fully sequenced *Vibrionaceae* (Pollack-Berti *et al*., 2010), so we wondered whether these homologs would act in a comparable manner in non-cholera vibrios despite the fact that these *Vibrio* species are often adapted to different environmental niches (Le Roux and Blokesch, 2018) and their homologous proteins might therefore serve different, adaptation-specific functions. We therefore compared the protein sequences of TfoX and TfoY from four different *Vibrio* species, *V. cholerae*, *V. parahaemolyticus*, *V. alginolyticus*, and *V. fischeri*, with the additional species chosen because previous studies had already described the genetic organization of their T6SS (Salomon *et al*., 2013; Salomon *et al*., 2014; Salomon *et al*., 2015; Speare *et al*., 2018). Moreover, these additional species have isolates that contain two T6SSs. We therefore considered the possibility that TfoX and TfoY, which are produced under vastly different conditions in *V. cholerae* (Metzger and Blokesch, 2016), might be specialized for the induction of a single T6SS in those organisms that harbor several systems. By aligning the protein sequences, we observed a high level of protein conservation for TfoX and TfoY, with a sequence identity of 68%/66%, 67%/66%, and 59%/67% when the homologous proteins of *V. parahaemolyticus* (TfoX*_Vp_*/TfoY*_Vp_*), *V. alginolyticus* (TfoX*_Va_*/TfoY*_Va_*), and *V. fischeri* (TfoX*_Vf_*/TfoY*_Vf_*) were compared to the respective homologs of *V. cholerae* (Fig. 1A). Given these identity values, we concluded that the TfoX of *V. parahaemolyticus* and *V. alginolyticus* were more closely related to the homologous protein of *V. cholerae*, whereas TfoX from *V. fischeri* diverged the most, which is in accordance with the phylogenetic distances between the species (Fig. 1A).

**Figure 1:**
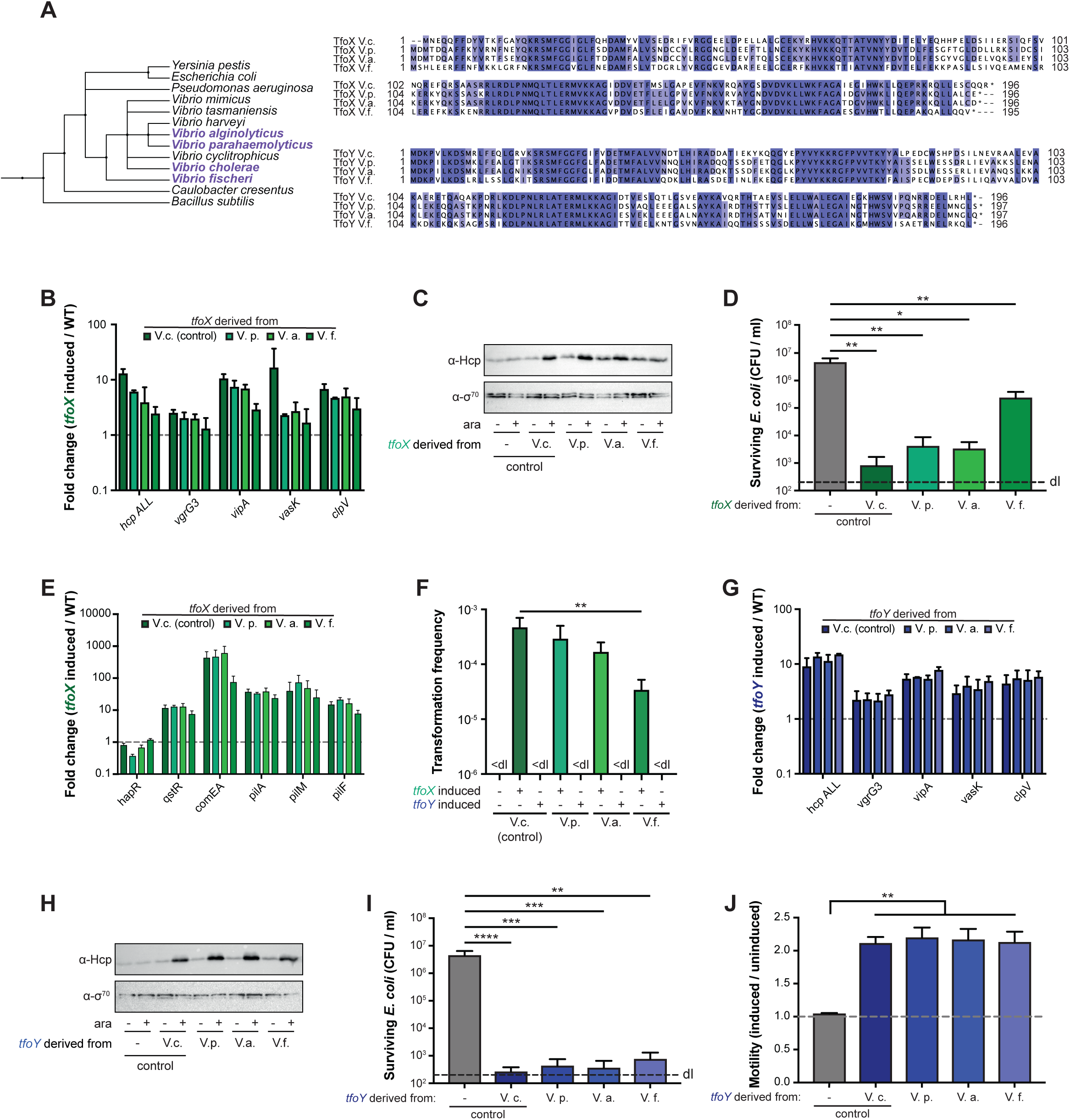
TfoX and TfoY proteins from diverse *Vibrio* species are functional in *V. cholerae*. (A) Phylogenetic tree and protein sequence conservation of TfoX and TfoY across *Vibrio* species. The phylogenetic tree was generated on phlyot.biobyte.de, a phylogenetic tree generator, based on NCBI taxonomy. The protein sequences of TfoX and TfoY were aligned using ClustalOmega, and subsequently modified with Jalview. Identical residues are highlighted in shades of blue (above threshold of 65%). (B-J) Chromosomally-located *tfoX* and *tfoY* from cholera (*V. cholerae*) and non-cholera *Vibrio* species (*V. parahaemolyticus*, *V. alginolyticus*, and *V. fischeri*) were expressed under the control of the arabinose-inducible P_BAD_ promoter in *V. cholerae*. The color code (TfoX-induced, green; TfoY-induced, blue) is used throughout the graphs. The WT strain lacking an inducible copy of *tfoX* or *tfoY* served as the negative control. The fold change (*tfoX*- or *foY*-induced over parental WT strain) of relative gene expression of (B, G) T6SS genes or (E) representative competence genes. (C, H) Detection of Hcp protein produced by TfoX- and TfoY-expressing cells. Cells were grown in the absence or the presence of the inducer (arabinose) as indicated below the image. Detection of σ70 served as a loading control. (D, I) Interspecies killing assay with *E. coli* as prey. *V. cholerae* harboring different inducible versions of (D) *tfoX* or (I) *tfoY* were co-cultured with the prey on LB agar plates supplemented with arabinose. The survival of the prey was determined on selective LB agar plates and is depicted as CFU per ml. (F) Natural transformation using genomic DNA is maintained in a *V. cholerae* strain expressing *tfoX* from non-cholera vibrios but that is non-functional upon *tfoY* expression. The indicated strains were grown under inducible conditions, and the genomic DNA of A1552-lacZ-Kan served as the transforming material. Transformation frequencies reflect the number of transformants divided by the total number of CFUs. < dl, below detection limit. (J) Motility was scored on soft agar with and without arabinose as an inducer. The motility phenotype was quantified as the ratio between the induced and uninduced conditions, as shown on the Y-axis. Abbreviations: V.c., *Vibrio cholerae* A1552, V.p., *Vibrioparahaemolyticus* RIMD2210633; V.a., *Vibrio alginolyticus* 12G01; V.f., *Vibrio fischeri* ES114. Bar plots represent the average of at least three independent biological replicates (± SD). Statistical significance is indicated (^∗^*p* < 0.05; ^∗∗^*p* < 0.01; ^∗∗∗^*p* < 0.001; ^∗∗∗∗^*p* < 0.0001).

Given the high level of protein conservation, we next tested whether the protein homologs from the non-cholera vibrios could function in *V. cholerae*. For this purpose, we cloned the respective genes preceded by an arabinose-inducible promoter (P_*BAD*_) onto a site-specific transposon (mini-Tn7) together with the gene that encodes the arabinose-responsive regulatory protein AraC (Table S1). Next, we grew the genetically engineered strains and the parental WT strain in the presence of arabinose (as inducer), isolated their RNA, and then quantified the transcript levels of representative T6SS-encoding genes. As shown in Fig. 1B, heterologous production of the diverse TfoX proteins resulted in increased T6SS transcript levels compared to the uninduced conditions, though the level of induction was slightly lower than the production of *V. cholerae*’s own TfoX, especially for the most diverging protein variant from *V. fischeri* (TfoX*_Vf_*). We confirmed these data at the protein level through the detection of the inner tube protein of the T6SS, Hcp (Fig. 1C). Consistent with the production of Hcp and the increase of the T6SS transcript level, these strains were able to kill *E. coli* in an interbacterial competition assay, with the least predatory behavior observed for the most divergent TfoX*_Vf_*-carrying variant (Fig. 1D).

TfoX of *V. cholerae* was first identified as the master regulator of natural competence for transformation (Meibom *et al*., 2005; Metzger and Blokesch, 2016). The TfoX-induced competence regulon includes those genes that encode the structural components of the DNA-uptake machinery, which primarily consists of a central pilus structure (encoded by *pil* genes; Seitz and Blokesch, 2013) and the DNA binding protein ComEA that is essential for reeling DNA across the outer membrane and into the periplasmic space (Seitz and Blokesch, 2014; Seitz *et al*., 2014; Matthey and Blokesch, 2016). We therefore tested whether the TfoX variants from the other *Vibrio* species would likewise foster competence-gene expression, which was indeed the case (Fig. 1E). Compared to the other variants, the more diverse *TfoX_Vf_* did not induce the same high transcript levels of *comEA*, which, accordingly, also led to lower transformation frequencies (Fig. 1F). Notably, *comEA* expression in *V. cholerae* is regulated by an intermediate transcription factor, QstR, which is dependent on the production of both TfoX and HapR (Lo Scrudato and Blokesch, 2013; Jaskólska *et al*., 2018). HapR is the master regulator of the QS system, meaning that *comEA* expression is dependent on a functional QS system and on high cell density. As *V. fischeri* - but not the other tested *Vibrio* species - lacks a QstR homolog, we suggest that TfoX_*Vf*_ induces *comEA* at lower levels, as it is adapted to the more diverse QS circuit of *V. fischeri* and the absence of QstR (Milton, 2006; Verma and Miyashiro, 2013).

We next tested the functionality of the TfoY homologs in *V. cholerae* by first testing the ability of the TfoY variants to foster natural transformation, given that these proteins are frequently annotated as “DNA transformation protein TfoX” or “TfoX-family DNA transformation protein” due to their similarity to TfoX. However, no transformants were detected upon the production of any of these proteins, indicating their inability to induce natural competence in *V. cholerae* (Fig. 1F). Conversely and consistent with the high sequence similarity of the proteins from the four organisms, all TfoY variants functioned at the same level as the *V. cholerae* TfoY to increase T6SS transcript levels (Fig. 1G), to induce Hcp protein production (Fig. 1H), and, consequently, to foster interbacterial killing of *E. coli* (Fig. 1I). In addition, the four TfoY variants were able to induce motility as a T6SS-independent phenotype, as visualized on swarming agar plates and quantified in Fig. 1J. Taken together, these data indicate that TfoX and TfoY are conserved proteins among these four *Vibrio* species and that the TfoY variants can fully replace the *V. cholerae* innate protein, while the TfoX variants are functionally replaceable at high levels for TfoX_*Vp*_/TfoX_*Va*_ and intermediate levels for TfoX*_Vf_*.

### TfoXbut not TfoYinduces V. fischeri’sprimary T6SS

While the complementation assay in *V. cholerae* described above provided a first insight into the conservation of TfoX and TfoY from diverse vibrios by indicating their similarity to those of *V. cholerae*, the experiments did not tell us whether these proteins would be involved in similar phenotypes as in *V. cholerae* in their native hosts. We therefore genetically engineered the different *Vibrio* species to place inducible copies of their own *tfoX* and *tfoY* genes onto a mini-Tn7 transposon that was then integrated into their own genomes (Table S1). Next, we assessed the different phenotypes under uninduced and induced conditions. For *V. fischeri* strain ES114 (Table S1), we showed that TfoX production led to a high induction of T6SS transcripts, while TfoY only partially induced a subset of T6SS genes (Fig. 2A). Consistent with these expression data, we witnessed the T6SS-mediated killing of *E. coli* only upon TfoX induction, while TfoY induction did not change the number of surviving co-cultured prey bacteria (Fig. 2B).

**Figure 2:**
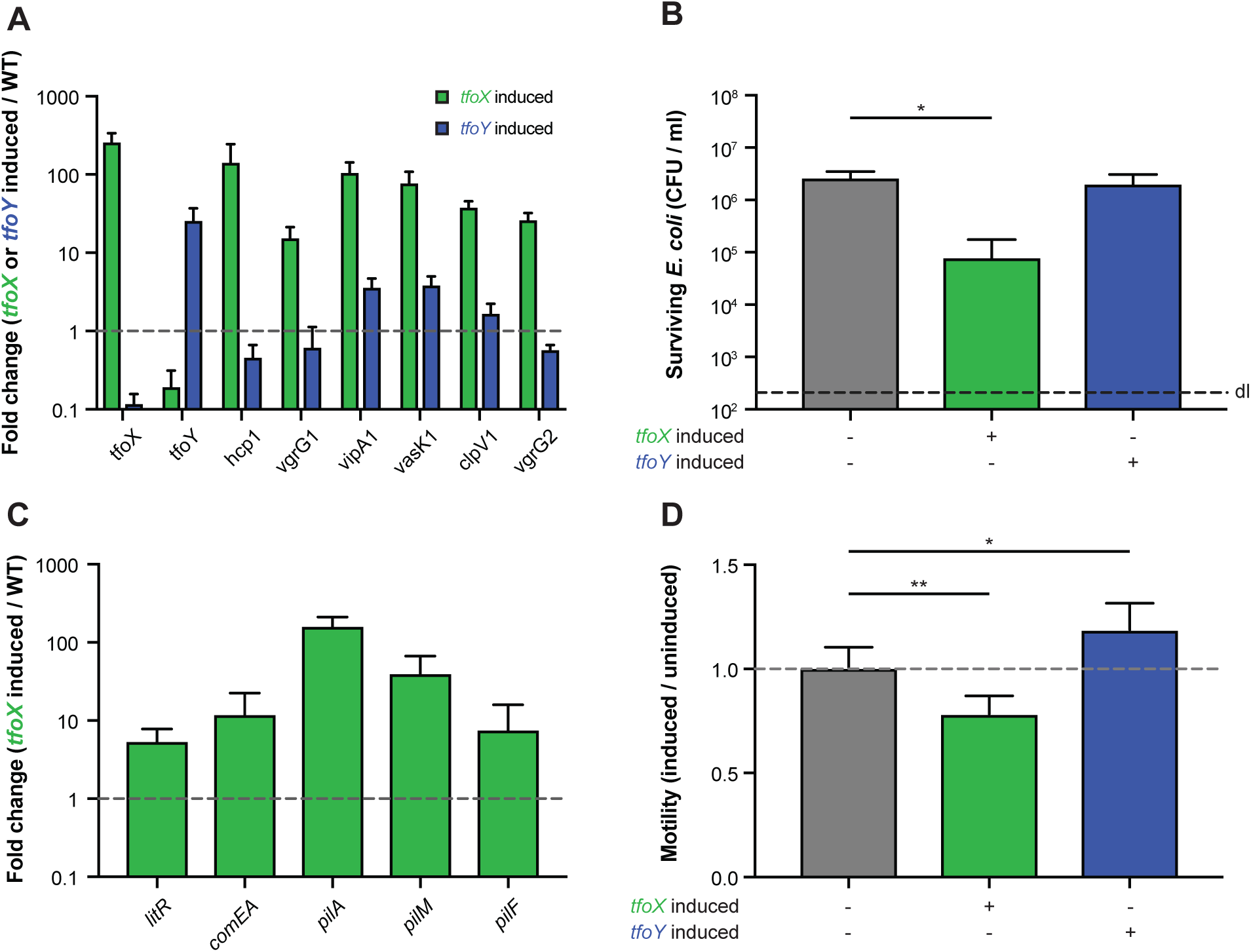
TfoX but not TfoY induce *V. fischeri’s* T6SS. The effect of TfoX (green) and TfoY (blue) production in *V. fischeri* ES114 was scored by qRT-PCR, and the relative expression values comparing uninduced versus induced conditions are indicated for representative (A) T6SS or (C) competence genes. (B) TfoX induction in *V. fischeri* leads to interbacterial killing of *E. coli* prey cells. *V. fischeri* strains were co-cultured with *E. coli* for 4 hours at 28°C on plain LBS agar plates supplemented with arabinose to induce the arabinose-inducible copy of *tfoX* or *tfoY* (as indicated below the graph). The values for the recovered prey are indicated on the Y-axis. (D) TfoY fosters motility in *V. fischeri*. Quantification of the motility phenotype of TfoX- or TfoY-induced bacteria normalized by the uninduced conditions. The WT strain without any inducible gene served as a control. (A-D) Bar plots represent the average of at least three independent biological replicates (± SD). Statistical significance is indicated (^∗^*p* < 0.05; ^∗∗^*p* < 0.01).

Pollak-Berti *et al*. showed that TfoX was required for natural competence for transformation in *V. fischeri* (Pollack-Berti *et al*., 2010), as is the case for *V. cholerae* (Meibom *et al*., 2005). We therefore measured relative transcript levels of representative competence genes in this organism and confirmed their induction upon TfoX production (Fig. 2C). Surprisingly, while we only observed a mild induction of *comEA*, we witnessed an almost 10-fold upregulation of *litR*, which encodes the main QS regulator LitR (homolog of *V. cholerae’s* HapR). This phenotype differs from what is known about competence regulation in *V. cholerae* in which case *hapR* transcript levels are left unchanged upon TfoX induction (Meibom *et al*., 2005; Metzger and Blokesch, 2016). We therefore speculate that an increase in LitR levels can compensate for the lack of the intermediate regulator QstR in *V. fischeri*. Indeed, the QstR of *V. cholerae* is required for competence-mediated DNA uptake in two ways: i) through direct and in-direct induction of certain competence genes, such as *comEA*, as described above; and ii) due to its ability to downregulate the *dns* gene (Lo Scrudato and Blokesch, 2013; Jaskólska *et al*., 2018) encoding for the extracellular nuclease Dns, which degrades transforming material outside the cell and within the periplasmic space (Blokesch and Schoolnik, 2008; Seitz and Blokesch, 2014). Notably, the HapR protein of *V. cholerae* also silences *dns* expression, but only partially in the absence of QstR (Lo Scrudato and Blokesch, 2013). It is therefore likely that the increased production of LitR upon TfoX production in *V. fischeri* compensates for the absence of QstR to ensure sufficient repression of *dns*, which, ultimately, would allow DNA uptake to occur. It is not clear, however, why the *comEA* transcripts were not induced to higher levels, though we recently demonstrated for *V. cholerae* that the *comEA* gene requires an additional but so far unidentified regulatory input apart from TfoX and QstR (Jaskólska *et al*., 2018). Indeed, we demonstrated that artificial QstR production was sufficient to induce the T6SS genes in this organism at comparable levels as the upstream regulatory protein TfoX, even in the absence of HapR, while the *comEA* transcript levels were only partially increased compared to a non-induced WT strain. We therefore hypothesize that this input signal might be conserved in *V. fischeri*, though missing under the tested conditions. This would lead to a high overall competence-gene induction upon TfoX production but lower *comEA* transcripts as compared to *V. cholerae* (Fig. 1E).

Lastly, we determined that the motility of *V. fischeri* changed upon TfoY production (Fig. 2D), even though the difference between the TfoY-induced and TfoY-uninduced state was not as pronounced as for *V. cholerae* (Fig. 1J). We argue that this might be caused by a basal production level of TfoY in this organism that does not rely on the artificial induction. A higher basal TfoY level would result in a higher basal motility, which would therefore appear as a less pronounced induction upon additional TfoY production. There was also a decrease in motility observable upon TfoX production (Fig. 1D), which is consistent with the idea of a basal TfoY level, as the expression data showed that TfoX significantly lowered the *tfoY* transcript levels (Fig. 2A). Likewise, TfoY production lowered *tfoX* transcript levels (Fig. 2A), suggesting that both proteins work independently of each other and are mutually exclusive in *V. fischeri*. This finding contradicts a previous study in which the authors concluded that TfoY was required for efficient TfoX-mediated transformation (Pollack-Berti *et al*., 2010). Notably, both studies used different conditions (e.g., inducible TfoX/TfoY expression in this study versus a transposon-inserted mutant of *tfoY* (Pollack-Berti *et al*., 2010)) and further studies are therefore required to conclusively show if TfoY is indeed involved in natural transformation in *V. fischeri* or not, as we suggest based on the herein-described data.

### TfoX and TfoY have a different impact on T6SS induction in V. alginolyticus

While *V. cholerae* strains and the examined *V. fischeri* strain ES114 contain only a single T6SS, several other *Vibrio* species contain more than one. We therefore asked whether, and if so how, TfoX and TfoY affect the production of different T6SSs within the same organism. To address this question, we first tested the contribution of these regulators to the two T6SSs (T6SS1 and T6SS2) of *V. alginolyticus* in an *E*. *coli*-killing assay using wildtype (WT) *V. alginolyticus* or its *hcp1* (referred to as ΔT6SS1 throughout the text), *hcp2* (referred to as ΔT6SS2 throughout the text), or double mutant as a predator. Consistent with a previous report by Salomon *et al*. (Salomon *et al*., 2015), WT *V. alginolyticus* did not show interbacterial predation at 37°C on standard LB medium (e.g., without additional salt; Fig. 3A). Upon production of TfoX or TfoY, however, predation by the WT was significantly increased. This effect was less pronounced but reproducible in the strain that only carried the complete T6SS2 gene set (ΔT6SS1), while a strain that only carried a complete T6SS1 gene set (ΔT6SS2) was not able to kill the *E. coli* cells, indicating that T6SS1 is non-functional at this temperature. In contrast, both systems were inducible at 30°C, with T6SS1 specifically induced by TfoY and not TfoX, while T6SS2 was responsive to both regulatory proteins (Fig. 3B). Basic killing activity was also observed in the WT and the mutant lacking T6SS1 but not in a mutant lacking T6SS2, suggesting that T6SS2 is slightly induced under the tested conditions but can be further boosted upon TfoX and TfoY production (Fig. 3B). While this finding seems at first glance to contradict a previous study that reported basal activity for T6SS1 (Salomon *et al*., 2015), it should be noted that the experimental setup differed in that we used an assay in which the *Vibrio* are tested at high cell density, as was previously developed for *V. cholerae* (Borgeaud *et al*., 2015), while Salomon *et al*. first diluted the cells to low densities. The rationale behind our experimental setup was the knowledge that the TfoX-mediated pathway requires co-induction by HapR in *V. cholerae* (homologs are LuxR in *V. alginolyticus* and OpaR in *V. parahaemolyticus*) (Lo Scrudato and Blokesch, 2012; Metzger and Blokesch, 2016; Jaskólska *et al*., 2018), which we assumed to be a conserved feature in other *Vibrio* species (see data below for *V. parahaemolyticus*).

**Figure 3:**
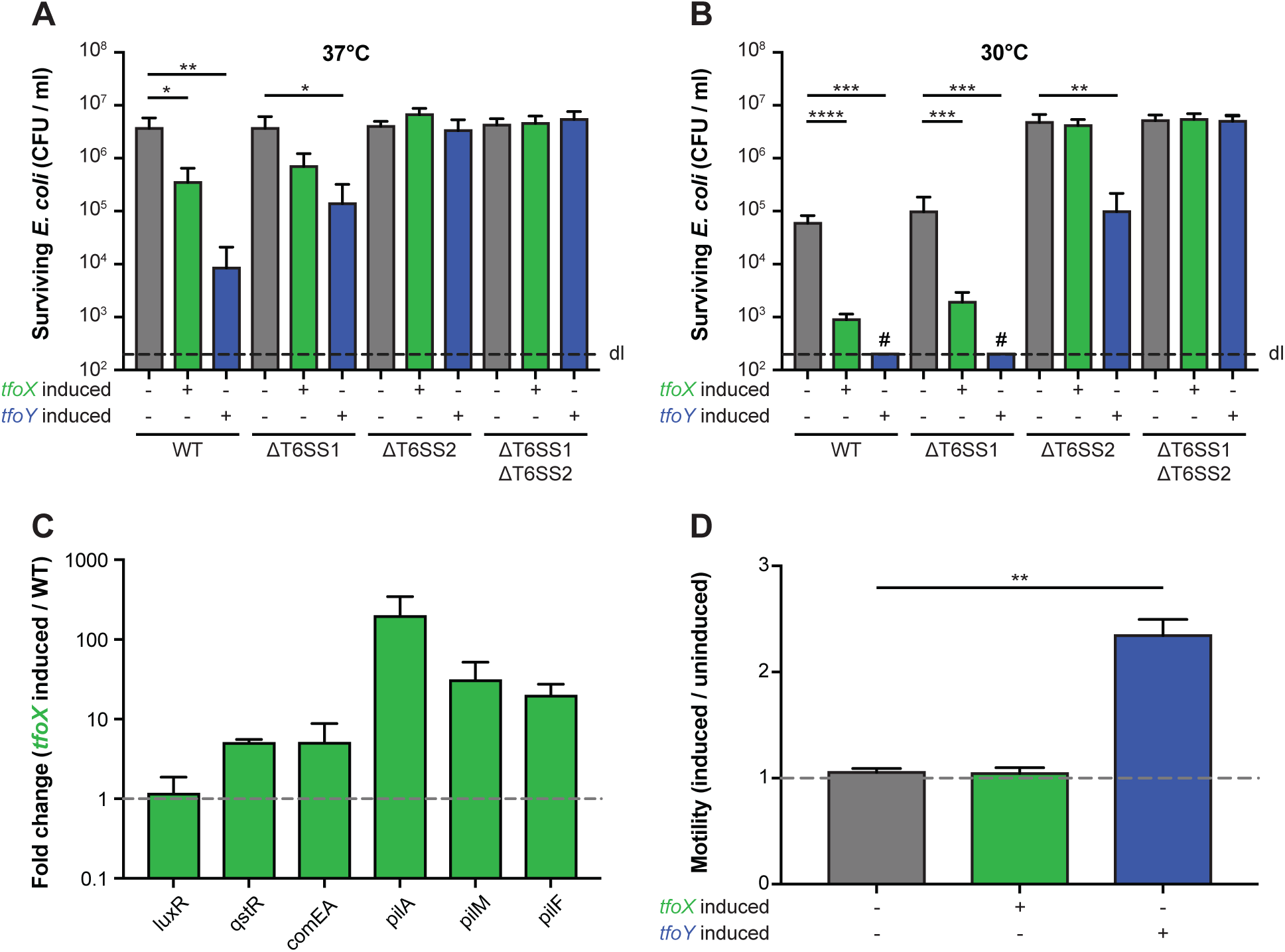
TfoY is a universal T6SS inducer in *V. alginolyticus*, while TfoX is specific for T6SS2. (A, B) Interspecies killing assay of *V. alginolyticus* with *E. coli* as prey. *V. alginolyticus* strains were co-cultured with *E. coli* prey for 4 hours at (A) 37°C (B) or 30°C on plain LB agar supplemented with arabinose to induce *tfoX* or *tfoY* (as indicated below the graph). The recovery of the prey is indicated as CFU/ml on the Y-axis. (C) TfoX-induces competence genes in *V. alginolyticus*. The relative expression of representative competence genes was scored in *tfoX*-induced and *tfoX*-uninduced strains, and the fold change between the values is indicated on the Y-axis. (D) Motility ratio between uninduced and induced *V. alginolyticus* strains is indicated. Details as in Fig. 2D. Abbreviations: WT, *V. alginolyticus* strain 12G01; ΔT6SS1, 12G01Δhcp1; ΔT6SS2, 12G01Δhcp2; ΔT6SS1ΔT6SS2, 12G01Δhcp1Δhcp2. Bar plots represent the average of at least three independent biological replicates (± SD). Statistical significance is indicated (^∗^*p* < 0.05; ^∗∗^*p* < 0.01; ^∗∗∗^*p* < 0.001; ^∗∗∗∗^*p* < 0.0001).

In addition to T6SS activation, we also tested the level of competence genes transcripts and bacterial motility upon TfoX and TfoY induction. As shown in Fig. 3C, a significant TfoX-mediated induction of representative competence genes occurred, indicating the protein’s conserved action. Notably, the *comEA* transcript induction levels again turned out to be unexpectedly low compared to the induction levels in *V. cholerae* (Fig. 1E). ComEA plays a primary role in ratcheting the DNA into the periplasmic space in competent bacteria (Seitz *et al*., 2014), and we therefore expect that these low transcript levels would lead to insufficient amounts of the ComEA proteins and defective DNA uptake. We speculate that the additional but so far unknown regulatory input required for *comEA* expression mentioned above (Jaskólska *et al*., 2018) might also be a prerequisite in *V. alginolyticus* and that this pathway might be non-functional in either the strain used in this study or under the tested conditions or both. The motility-inducing phenotype of TfoY, on the hand, was very pronounced in *V. alginolyticus*, strengthening the idea that TfoY protein levels and a high motility are linked throughout the genus *Vibrio*, even in organisms that contain lateral flagella in addition to the polar flagellum (McCarter, 2001).

### TfoX and TfoY are both dedicated to one specific T6SS in V. parahaemolyticus

Lastly, we tested the contribution of TfoX and TfoY to T6SS regulation in *V. parahaemolyticus*. As for *V. alginolyticus*, this bacterium also contains two T6SSs, T6SS1 and T6SS2, which were previously shown to be differentially regulated from each other with respect to temperature, salinity, QS inputs and surface sensing (Salomon *et al*., 2013). Indeed, T6SS1 of *V. parahaemolyticus* was found to be most active under marine-like conditions and at low cell densities while system 2 was more adapted to lower salt conditions (e.g., LB medium) and higher cell densities (Salomon *et al*., 2013). Notably, these data on T6SS2 regulation and a previous study that suggested T6SS2 was involved in adhesion to host cells were solely based on the Hcp2 protein levels (inside and outside the cells), while interbacterial killing activity was not directly observed for T6SS2 (Yu *et al*., 2012; Salomon *et al*., 2013). Indeed, the QS-dependent T6SS2 regulation of Hcp production confirmed previous results in *V. cholerae* (Ishikawa *et al*., 2009). However, despite the fact that pandemic *V. cholerae* produce detectable levels of Hcp at high cell densities *in vitro*, the bacteria are unable to kill prey under such conditions (Pukatzki *et al*., 2006), which can be overcome by TfoX or TfoY production (Metzger *et al*., 2016). Indeed, we previously demonstrated that upon induction of these two regulators, the Hcp levels increased further, and T6SS-mediated prey killing was observed. Notably, TfoY-mediated T6SS activity in *V. cholerae* occurred independently of HapR, indicating that TfoY acts independently from the QS pathway, in contrast with TfoX-mediated T6SS activation (Borgeaud *et al*., 2015; Metzger *et al*., 2016).

In *V. parahaemolyticus*, both TfoX and TfoY production led to significant *E. coli* killing (Fig. 4A). Interestingly, when testing *V. parahaemolyticus* strains that contained only a complete T6SS2 gene set (ΔT6SS1) or only all T6SS1 genes (ΔT6SS2), it was clear that TfoX specifically induced T6SS2, while TfoY solely led to T6SS1-mediated prey killing (Fig. 4A). These data also suggested that the T6SS2 was slightly active in both the WT and the strain lacking T6SS1, even without TfoX/TfoY production. We then determined that low levels of TfoX or TfoY did not cause this basal activity, as deletion of the indigenous genes did not abolish it (Fig. 4B). Interestingly, a recent study showed T6SS2-dependent killing of a *V. parahaemolyticus* strain that lacked a newly identified effector immunity pair (RhsP and RhsPi) by its parental WT strain, even in the absence of TfoX or TfoY induction (Jiang *et al*., 2018). These data therefore suggest that the basal activity that we observed against *E. coli* as a prey in this study might be much more enhanced in killing non-immune siblings compared to other species or, alternatively, that the experimental conditions were different to the current work and that under such conditions the T6SS2 is highly active. Lastly, we cannot exclude the possibility that strain differences or the domestication of certain strains has caused the difference in basal T6SS activity. Notably, the focus of the current study was on the contribution of TfoX or TfoY to T6SS activity, which we unambiguously demonstrate to enhance T6SS activity in *V. parahaemolyticus*.

**Figure 4:**
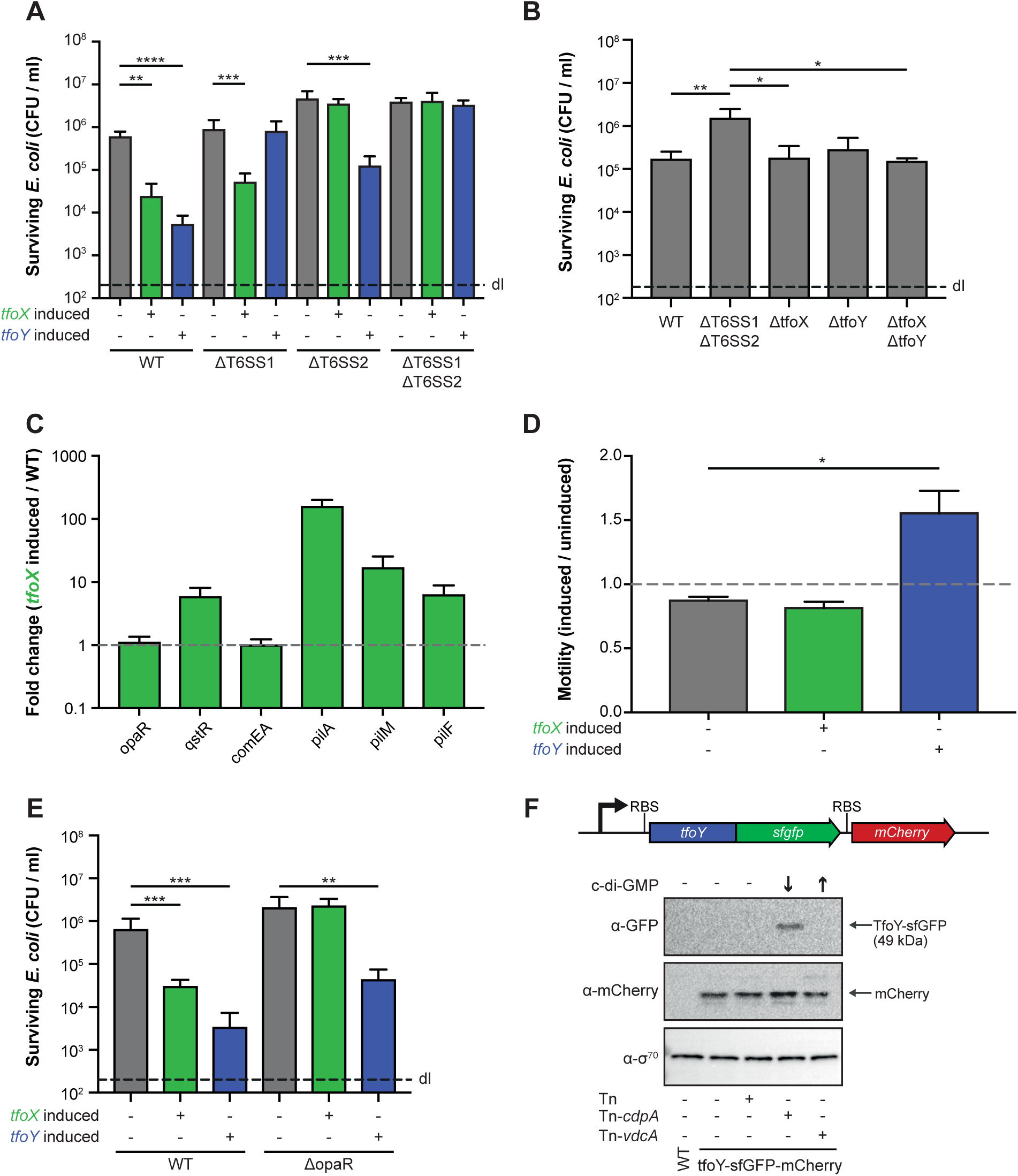
TfoX and TfoY each induce a dedicated T6SS in *V. parahaemolyticus*. (A, E) Interspecies killing of *E. coli* by *V. parahaemolyticus. V. parahaemolyticus* strains (WT, T6SS1-, T6SS2-, T6SS1/T6SS2-minus or opaR-minus) were co-cultured with *E. coli* prey for 4 hours at 30°C on LB agar plates supplemented with arabinose to induce *tfoX* or *tfoY* (as indicated below the graph). The recovery of the prey is indicated on the Y-axis. (B) Basal T6SS activity is independent of TfoX and TfoY. WT *V. parahaemolyticus* or its *tfoX*-, *tfoY*-, or *tfoX/tfoY*-minus variants were tested for their ability to reduce the number of *E. coli* prey cells. The T6SS double mutant (ΔT6SS1ΔT6SS2) served as a control. (C) Fold change (*tfoX*-induced over the WT parental strain) of relative gene expression of representative competence genes. (D) Quantification of TfoX- or TfoY-induced motility phenotypes. Details as in Fig. 2D. (F) TfoY is produced under low c-di-GMP conditions in *V. parahaemolyticus*. Detection of TfoY-sfGFP and mCherry by western blotting in *cdpA*- or *vdcA*-inducible reporter strains that contain increased or decreased intracellular c-di-GMP levels, as indicated by the arrows. The transposon-less or empty transposon-carrying reporter strain as well as the parental WT strain served as controls. Detection of σ70 served as a loading control. RBS, ribosome-binding site. Abbreviations: WT, *V. parahaemolyticus* POR1 (RIMD2210633 derivative); ΔT6SS1, POR1Δhcp1; ΔT6SS2, POR1Δhcp2; ΔT6SS1ΔT6SS2, POR1Δhcp1Δhcp2. Bar plots represent the average of at least three independent biological replicates (± SD). Statistical significance is indicated (^∗^*p* < 0.05; ^∗∗^*p* < 0.01; ^∗∗∗^*p* < 0.001; ^∗∗∗∗^*p* < 0.0001).

Examining other TfoX- and TfoY-mediated phenotypes confirmed the increase of competence gene transcripts and enhanced motility, respectively (Fig. 4C and D). However, the *comEA* levels were again unexpectedly low, which explains why we were unable to naturally transform *V. parahaemolyticus* even after TfoX induction (data not shown). The induction levels of the gene encoding *qstR* were comparable with what we observed for *V. cholerae* (Fig. 1E) where TfoX-dependent T6SS induction seemed functional (Fig. 4A), so we hypothesize that the transformation defect was *comEA*-specific, as discussed above, and not based on a general mutation within the QS cascade. Previous studies in different *Vibrio* species described gain-of-function mutations in LuxO, a repressor of HapR/LitR/OpaR synthesis (Gode-Potratz and McCarter, 2011; Kimbrough and Stabb, 2015; Stutzmann and Blokesch, 2016) and clinical *V. parahaemolyticus* isolates with such *luxO* mutantions were previously described to be locked in a pathogenic state (Kernell Burke *et al*., 2015). To exclude that this has happened in the strain we were working with, we first sequenced *luxO* and *opaR* and confirmed that both genes were mutation free. Additionally, we deleted *opaR* from the WT strain and compared this strain to its TfoX- or TfoY-inducible derivatives (Table S1) in an *E*. *coli*-killing assay. As shown in Fig. 4E, the TfoY-induced prey killing was maintained upon deletion of *opaR*, while the TfoX-mediated T6SS activity was abrogated. Since OpaR is produced at high cell density, its deletion blocks QS-dependent signalling. We therefore conclude that the WT strain has a functional OpaR and that TfoX-mediated T6SS induction in *V. parahaemolyticus* requires co-regulation by OpaR, similar to *V. cholerae* (Borgeaud *et al*., 2015; Metzger *et al*., 2016).

Our data suggest that TfoX induces the T6SS2 in *V. parahaemolyticus*. As previous studies showed that the chitin sensors ChiS, TfoS, and the downstream-regulated small RNA TfoR, which is required for *tfoX* mRNA translation, are highly conserved in diverse *Vibrio* species (Li and Roseman, 2004; Yamamoto *et al*., 2011; Yamamoto *et al*., 2014), we suggest that TfoX is exquisitely produced upon growth on chitinous surfaces, as first demonstrated for *V. cholerae* (Meibom *et al*., 2004; Meibom *et al*., 2005). The production of TfoY is less understood. In a previous study on *V. cholerae*, we showed that lowered c-di-GMP levels led to the production of TfoY under standard laboratory conditions (e.g., in LB medium) and we and others suggested that this c-di-GMP-dependent control occurred at the translational level (Inuzuka *et al*., 2016; Metzger *et al*., 2016). Interestingly, Gode-Potratz *et al*. demonstrated that growth on surfaces lowered the levels of the secondary messenger c-di-GMP in *V. parahaemolyticus* (Gode-Potratz *et al*., 2011). Consistently, Salomon *et al*. established that surface sensing correlated with Hcp1 production in this organism (Salomon *et al*., 2013). Based on these data, we speculated that surface sensing, TfoY production, and T6SS1 induction might be linked. To address this hypothesis and especially the link between low c-d-GMP levels and TfoY production, we genetically engineered *V. parahaemolyticus* to carry a translational fusion between its indigenous *tfoY* gene and the gene that encodes super-folder GFP (sfGFP). In addition, we included a transcriptional mCherry-encoding reporter gene behind this construct, which, ultimately, allowed us to monitor the transcriptional and translational control of *tfoY*. Next, we incorporated either an empty mini-Tn7 transposon (Tn) or transposons that carried arabinose-inducible genes coding for a c-di-GMP-producing diguanylate cyclase (Tn-*vdcA*) or for a phosphodiesterase (Tn-*cdpA*) into this strain (Table S1). These strains as well as the transposon-deficient parental strain and the WT without any fluorescent protein-encoding genes were then grown in the presence of the inducer arabinose followed by a western blot analysis to visualize the TfoY-sfGFP fusion protein or the transcriptional reporter protein mCherry. Using this approach, we observed that TfoY was produced solely at low ci-di-GMP levels (Fig. 4F). Importantly, based on the comparable levels of mCherry in all reporter strains, we concluded that the *tfoY* transcript levels did not significantly change in response to increased or decreased c-di-GMP levels (Fig. 4F). This finding contradicted a recent study in *V. cholerae* that concluded that TfoY was induced at both low and high intracellular concentrations of c-di-GMP and that this regulation occurred at the translational and transcriptional levels, the latter due to activation by the c-di-GMP-dependent transcription factor VpsR (Pursley *et al*., 2018). We therefore constructed a similar dual translational-transcriptional reporter strain in *V. cholerae* (Table S1) and tested this strain under normal, increased, or decreased c-di-GMP levels. Using this approach, we were able to detect an increase in the TfoY protein under low c-di-GMP levels compared to normal conditions, while an increase in intracellular c-di-GMP did not result in higher TfoY protein levels (Fig. S2), consistent with our previous findings (Metzger *et al*., 2016). Importantly, we also did not observe a change in the transcriptional reporter under high c-di-GMP conditions (Fig. S2), despite normal VpsR function in the pandemic *V. cholerae* strain (A1552) used in this study (Yildiz *et al*., 2001), which confirmed the data presented above for *V. parahaemolyticus*. An explanation for this discrepancy might be that we engineered the construct at the native locus of *tfoY* within the *V. parahaemolyticus* or *V. cholerae* genomes, while Pursley *et al*. were unsuccessful in detecting tagged TfoY variants using a similar approach. They therefore decided to construct a TfoY-GFP translational-fusion-encoding reporter gene on a plasmid, which could change the expression pattern compared to the gene’s native locus. In addition, these authors performed their experiments in a *Vibrio* polysaccharide mutant as parental strain (ΔvpsL), which is deficient in biofilm formation (Pursley *et al*., 2018), while our strain maintained a normal ability for this process.

Collectively, we conclude that the lowered c-di-GMP levels observed in surface-grown *V. parahaemolyticus* (Gode-Potratz *et al*., 2011) might trigger TfoY induction, which, subsequently, induces the T6SS1 and subsequently initiates motility, though the exact mechanism for the latter remains to be discovered. The data provided here for *V. parahaemolyticus* therefore support our previous suggestion that TfoY triggers a defensive escape reaction that might allow *Vibrio* species to defend themselves against bacterial competitors and eukaryotic predators. Interestingly, a recent study that profiled the gene expression of *V. parahaemolyticus* reported increased *tfoY* expression (VP1028; falsely annotated as *tfoX* and therefore discussed as competence regulator in this study) within infected infant rabbits when compared to *in vitro* conditions (Livny *et al*., 2014), suggesting that TfoY might also play a role *in vivo* and therefore in pathogenesis.

## Conclusion

In this study, we demonstrated that the link between the TfoX and TfoY regulatory proteins and T6SSs is highly conserved among cholera and non-cholera *Vibrio* strains (Fig. 5), highlighting the general importance of these regulators. Interestingly, TfoX induced at least one T6SS in every tested species. Hence, we suggest that chitin-mediated TfoX production (Meibom *et al*., 2005; Metzger and Blokesch, 2016) and the resulting coupling between T6SS-facilitated neighbor killing and competence-mediated DNA uptake (Borgeaud *et al*., 2015) might be an ancient phenotype maintained in most *Vibrio* species. In contrast, the effect of TfoY on T6SS turned out to be more versatile (Fig. 5). While the c-di-GMP-dependent inhibition of TfoY translation seemed conserved, at least in the two tested *Vibrio* species (Fig. 4E and Fig. S2), the regulons seems to have diverged in the different organisms. Indeed, while TfoY induces both T6SSs in *V. alginolyticus* and one T6SS in *V. cholerae* and *V. parahaemolyticus* (out of two T6SSs present in the latter bacterium), this regulator does not impact the activity of the T6SS of *V. fischeri* under the tested conditions. Interestingly, several isolates of *V. fischeri*, other than the squid isolate ES114, contain a secondary T6SS. This T6SS2 is located on a strain-specific genomic island on the small chromosome and was recently shown to contribute to competitor elimination within the host’s light organ (Speare *et al*., 2018). While we were unable to genetically engineer the fish symbiont MJ11 strain for technical reasons, it is tempting to speculate that this secondary T6SS2 of *V. fischeri* might be controlled by the TfoY protein. Notably, Speare *et al*. proposed the possibility that this secondary T6SS2 was present in the common ancestor of *V. fischeri* and related *Vibrio* species but was otherwise lost in niche-adapted strains for which interbacterial killing is no longer advantageous. We extend this hypothesis and speculate that niche-adapted vibrios might no longer require a TfoY-mediated defensive response against either competing bacteria or grazing predators. Future studies are required to address these interesting evolutionary hypotheses.

**Figure 5:**
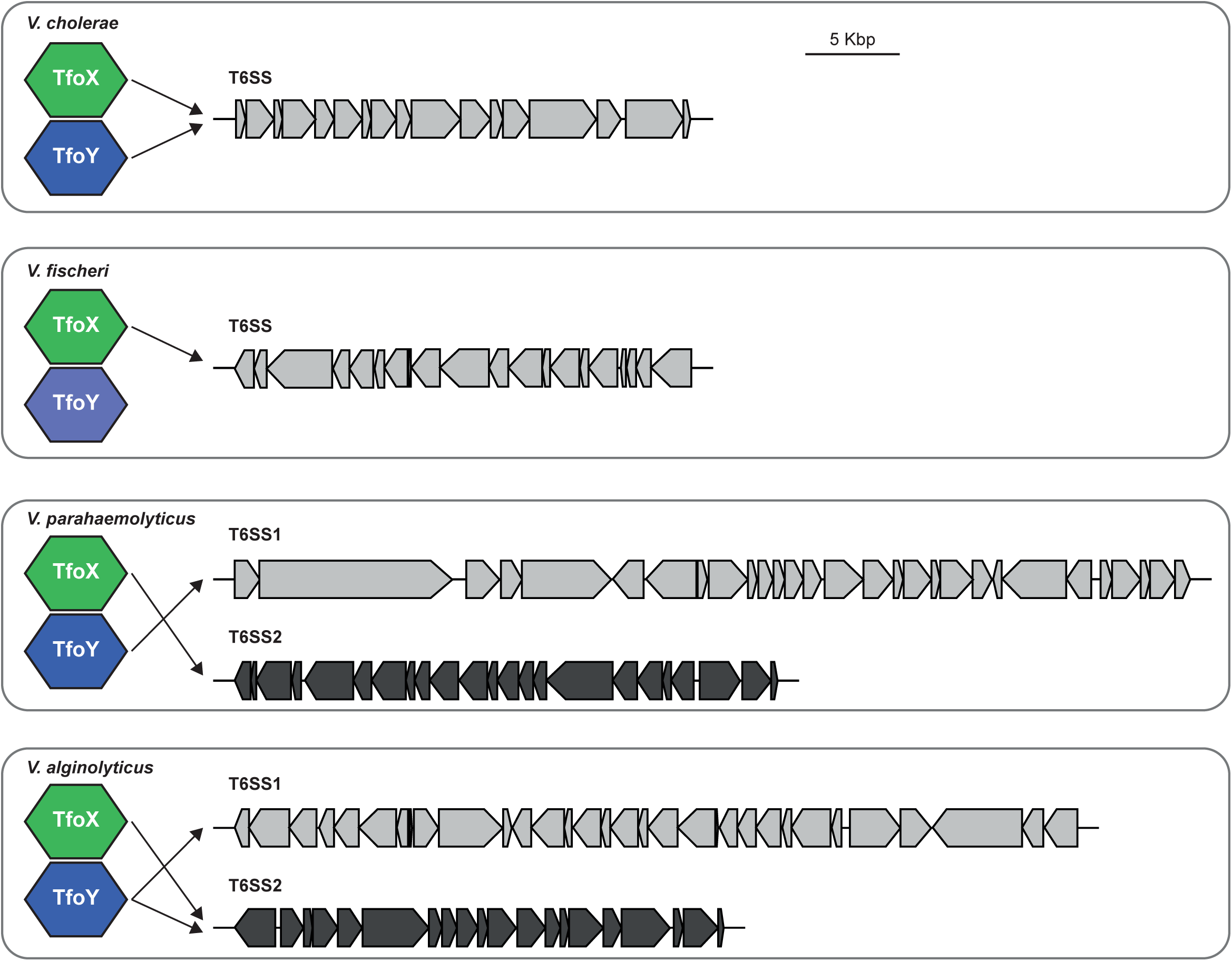
Summary scheme of TfoX- and TfoY-mediated T6SS induction in cholera and non-cholera *Vibrio* species. The T6SS clusters of *V. cholerae, V. fischeri*, *V. alginolyticus*, and *V. parahaemolyticus* are indicated (scale corresponds to 5 kbp as indicated). Arrows indicate positive regulation by the respective regulator TfoX or TfoY.

## Experimental procedures

### Bacterial strains, plasmids, and growth conditions

The *V. cholerae*, *V. alginolyticus*, *V. parahaemolyticus*, and *V. fischeri* strains and plasmids used in this study are listed in Table S1. *Escherichia coli* strains DH5α (Yanisch-Perron *et al*., 1985), TOP10 (Invitrogen), SM10λpir (Simon *et al*., 1983), S17-1λpir (Simon *et al*., 1983) and MFDpir (Ferrieres *et al*., 2010) were used for cloning purposes and/or served as donor in bacterial mating experiments.

Unless otherwise stated, the *V. cholerae*, *V. alginolyticus*, *V. parahaemolyticus*, and *E. coli* strains were grown aerobically in Lysogeny broth (LB; 10 g/L of tryptone, 5 g/L of yeast extract, 10 g/L of sodium chloride; Carl Roth) or on LB agar plates at 30°C or 37°C. *V. fischeri* strains were cultured aerobically in LB salt (LBS) medium (10 g/L of tryptone, 5 g/L of yeast extract, 20 g/L of sodium chloride, 20 mM Tris-HCl [pH 7.5], 0.2% glycerol, adapted from (Dunlap, 1989; Graf *et al*., 1994)) or LBS agar plates at 28°C. LB/LBS motility plates had less agar (0.3%) than standard LB/LBS agar plates (1.5%). The following were added if required at the given concentrations: arabinose (0.02% or 0.2%), diaminopimelic acid (DAP; 0.3 mM), or the antibiotics kanamycin (75 μg/ml), gentamicin (50 μg/ml), and chloramphenicol (2.5 μg/ml or 5 μg/ml). L-arabinose-supplemented medium was used for the expression of *tfoX* or *tfoY* under the control of the P_BAD_ promoter. DAP was added as an essential growth supplement for *E. coli* strain MFDpir (Ferrieres *et al*., 2010). Medium without DAP was used to counter-select MFDpir strains after tri-parental mating with *V. fischeri*. For the *E. coli* counter-selection after tri-parental mating with *V. cholerae, V. alginolyticus*, and *V. parahaemolyticus*, Thiosulfate Citrate Bile Salts Sucrose (TCBS) agar plates were used and prepared following the manufacturer’s instructions (Sigma-Aldrich/Fluka, Buchs, Switzerland). Marine broth 2216 (BD Difco^™^ 2216) was used to isolate single colonies of *V. alginolyticus* and *V. parahaemolyticus* after *E. coli* counter-selection.

### Genetic engineering of strains and plasmids

DNA manipulations were performed according to standard molecular biology-based protocols (Sambrook *et al*., 1982). Enzymes were purchased from the listed companies and were used as recommended by the manufacturer: Pwo polymerase (Roche), Taq polymerase (Promega), restriction enzymes (New England Biolabs). Following initial screening by PCR (using bacterial cells as templates), genetically engineered strains and plasmids were verified by Sanger sequencing (Microsynth, Switzerland).

*V. cholerae* strains were genetically modified using both a gene-disruption method based on the counter-selectable plasmid pGP704-Sac28 (Meibom *et al*., 2004) or the TransFLP gene disruption method previously described by our group (De Souza Silva and Blokesch, 2010; Blokesch, 2012b; Borgeaud and Blokesch, 2013). This transformation-based genetic engineering technique was used to replace the *tfoY* gene by a translational and transcriptional *tfoY*-*mCherry::f* fusion at the natural locus on the *V. cholerae* A1552 chromosome. Plasmid pGP704-Sac-Kan (see below), derived from pGP704-Sac28, was used for genetic modifications of *V. parahaemolyticus*.

To construct the counter-selectable plasmid pGP704-Sac-Kan, the *aph* gene was amplified from a kanamycin-resistance gene (*aph*)-carrying transposon using primers with overhanging *SspI* and *BsaI* restriction sites. A restriction digestion was performed on both the PCR product and the vector pGP704-Sac28, which were subsequently ligated, to replace the *bla* gene of pGP704-Sac28 with the *aph* gene, resulting in plasmid pGP704-Sac-Kan (Table S1).

All pBAD-derived plasmids harboring *tfoX*-*strep* or *tfoY*-*strep* genes from different *Vibrio* species were constructed in the following way: The genes were amplified with *StreptagII*-encoding primers using gDNA of the respective parental *Vibrio* strain as a template. The restriction-enzyme-digested PCR product was subsequently cloned into plasmid pBAD/MycHisA (Table S1). The resulting plasmids served as templates for fragments containing *araC*, the arabinose-inducible promoter PBAD, and the given *tfoX*, *tfoY* gene, which were subcloned into the mini-Tn7-containing delivery plasmids (Table S1). The plasmid pGP704-Tn-Cm^R^ is a derivative of pGP704-mTn7-minus-SacI (Nielsen *et al*., 2006), where the *aacC1* gene within the mini-Tn7 was replaced by the *cat* gene. To this end, the PCR-amplified *cat* gene (including its promoter) of pBR-FRT-Cat-FRT2 (Metzger *et al*., 2016) was cloned into pGP704-mTn7-minus-SacI, which was digested using restriction enzymes *SbfI* and EcoRV. For the insertion of this mini-Tn7 transposon into the *Vibrio spp*. chromosomes, a tri-parental mating strategy was employed (Bao *et al*., 1991). The donor plasmids are indicated in Table S1.

### Natural transformation assay

Natural transformation assays in liquid were performed with minor modifications to the previous protocol (Lo Scrudato and Blokesch, 2012). This assay is chitin-independent and uses strains carrying an arabinose-inducible copy of *tfoX*-*strep* or *tfoY*-*strep* (both from *V. cholerae*, *V. alginolyticus*, *V. parahaemolyticus*, or *V. fischeri*) on the chromosome. *V. cholerae* strains were pre-grown overnight in LB medium and further grown in the presence of the arabinose inducer up to an optical density at 600 nm (OD_600_) of 1.0. At this point, 0.5 ml of the cultures were supplemented with 1 μg of genomic DNA. For all natural transformation assays, the genomic DNA of A1552-lacZ-Kan (Marvig and Blokesch, 2010) served as transforming material. Cells were further incubated under shaking conditions at 30°C for 4 hours. Serial dilutions were spotted on LB to count the total number of cells and on kanamycin-containing LB agar plates to select the transformants. Transformation frequencies were calculated as the number of colony-forming units (CFUs) of the transformants divided by the total number of CFUs. Averages of at least three biologically independent experiments are provided. For statistical analyses, the data were log-transformed and significant differences were determined by the two-tailed Student’s t-test. When no transformants were recovered, the value was set to the detection limit to allow for statistical analysis.

### Gene expression analysis by quantitative reverse transcription PCR (qRT-PCR)

Quantitative reverse transcription PCR (qRT-PCR)-based transcript scoring in *V. cholerae*, *V. alginolyticus*, *V. parahaemolyticus*, and *V. fischeri* was performed following a previously established protocol (Lo Scrudato and Blokesch, 2012). Strains with and without an arabinose-inducible copy of specified genes (*tfoX*-*strep* or *tfoY*-*strep*) were grown for 6 hours in 2.5 ml LB or LBS supplemented with 0.2% arabinose. Cultures (2 ml) were processed for RNA isolation and subsequent cDNA synthesis. Relative gene expression values were normalized against the *gyrA* transcript levels. Fold changes were determined using the relative expression values of the induced strains divided by the values of the parental WT strain (as specified for each inducible construct in the figures). Averages of at least three biologically independent experiments (± standard deviation) are provided.

### Interbacterial killing assay using E. coli as prey

The *E. coli* killing assay was performed following a previously established protocol with minor adaptions (Borgeaud *et al*., 2015). The *E. coli* prey cells and the given predator cells were mixed at a ratio of 10:1 and spotted onto membrane filters on pre-warmed LB and/or LBS agar plates (± 0.2% ara). After 4 h of incubation at 28°C (predator: *V. fischeri)* or 30°C and 37°C (as indicated for the predators *V. cholerae*, *V. alginolyticus*, *and V. parahaemolyticus*), bacteria were resuspended and serial dilutions were spotted onto antibiotic-containing LB agar plates to enumerate the CFUs (shown as CFU/ml). Arabinose-uptake-deficient *E. coli* TOP10 and its derivative TOP10-TnKan served as prey. Significant differences were determined by a two-tailed Student’s t-test on log-transformed data of at least three biological replicates. If no prey cells were recovered, the value was set to the detection limit to allow statistical analysis.

### Motility assay

The motility of the *Vibrio* species was assessed by spotting 2 μl of the respective overnight culture onto freshly prepared LB and/or LBS motility agar plates (containing 0.3% agar) with or without 0.2% arabinose. Following the incubation at 30°C for 6h for *V. cholerae*, *V. parahaemolyticus*, and *V. alginolyticus*, or at room temperature for 7h for *V. fischeri*, the diameters of the bacterial swarming were determined. The motility induction was calculated by dividing the swarming diameter of induced versus uninduced strains. The averages of at least three independent experiments (± standard deviation) are provided. For statistical analyses, a two-tailed Student’s t-test was performed.

### SDS-PAGE and western blotting

Cell lysates were prepared as described previously (Metzger *et al*., 2016). In brief, after cultivation with or without arabinose for 3 or 6 hours, bacterial cell pellets were resuspended in Laemmli buffer, adjusting for the total number of bacteria according to the OD_600_ values. Proteins were separated by sodium dodecyl sulfate (SDS)-polyacrylamide gel electrophoresis and western blotted as described (Lo Scrudato and Blokesch, 2012). Primary antibodies against Hcp (Eurogentec; (Metzger *et al*., 2016)), GFP (Roche, Switzerland), and mCherry (BioVision, USA distributed via LubioScience, Switzerland) were used at 1:5,000 dilutions, and *E. coli* Sigma70 (BioLegend, USA distributed via Brunschwig, Switzerland) was used at a 1:10,000 dilution. Goat anti-rabbit horseradish peroxidase (HRP) and goat anti-mouse HRP (both diluted 1:20,000; Sigma-Aldrich, Switzerland) served as secondary antibodies. Lumi-Light^PLUS^ western blotting substrate (Roche, Switzerland) was used as an HRP substrate and the signals were detected using a ChemiDoc XRS+ station (BioRad).

## Acknowledgments

We thank members of the Blokesch lab and Frédérique Le Roux for fruitful discussions, Nina Vesel for constructing plasmid pGP704-Sac-Kan, and Sandrine Stutzmann, Julien Chambaud, and Tiziana Scrignari for technical assistance. We acknowledge Dor Salomon for providing the *V. parahaemolyticus* and *V. alginolyticus* strains, for sharing unpublished data, and for arranging co-submission. This work was supported by the Swiss National Science Foundation (31003A 162551) and a Consolidator Grant from the European Research Council (ERC; 724630-CholeraIndex). M.B. is a Howard Hughes Medical Institute (HHMI) International Research Scholar (grant 55008726).

## Author contributions

Conception, design and analysis: L.C.M, N.M., and M.B.; performed research: L.C.M, N.M., C.S., E.J.C., and M.B; wrote the manuscript: L.C.M. and M.B. with input from N.M.

## Supporting Information

Additional Supporting Information can be found in the online version of this article at the publisher’s webpage: XXX

**Table S1: Bacterial strains and plasmids used in this study**.

## Supporting figure legends

**Figure S1: TfoX- and TfoY-induced T6SS production is conserved in pandemic *V. cholerae* strains**.

Interspecies killing assay between diverse *V. cholerae* strains and *E. coli* as prey. The *V. cholerae* O1 El Tor strains tested are as follows: A1552, N16961rep (with repaired frameshift mutation in *hapR*), C6709, E7946, and P27459 as indicated below the graph. Co-culturing with *E. coli* occurred on LB agar plates supplemented with arabinose to induce *tfoX* or *tfoY* where indicated. The parental strains without inducible copies of the regulatory genes served as a control. Prey recovery is indicated as CFU/ml on the Y-axis. Bar plots represent the average of three independent biological replicates (± SD). Statistical significance is indicated (^∗^ < 0.05; ^∗∗^*p* < 0.01; ^∗∗∗^*p* < 0.001; ^∗∗∗∗^*p* < 0.0001).

**Figure S2: TfoY production is translationally but not transcriptionally controlled by c-di-GMP in *V. cholerae***.

*V. cholerae* reporter strains carrying a gene encoding for a translational fusion between TfoY and mCherry (*tfoY*-*mCherry*) with or without a downstream transcriptional reporter gene (*gfp*) at the gene’s native chromosomal locus were genetically manipulated to insert inducible copies *vdcA* or *cdpA* into their genome (inside a mini-Tn7 transposon). Cells were then grown under inducible conditions to increase or decrease intracellular c-di-GMP concentrations, as shown by the arrows above the images. Detection of TfoY-mCherry and GFP occurred through western blotting. The reporter strain lacking the transposon as well as the parental WT served as controls. Detection of o70 served as a loading control.

